# Cholesterol Oxidation Modulates the Formation of Liquid-Ordered Domains in Model Membranes

**DOI:** 10.1101/2021.05.24.445501

**Authors:** Paul Smith, Peter G. Petrov, Christian D. Lorenz

## Abstract

7-ketocholesterol (KChol) is one of the most cytotoxic oxysterols found in the plasma membrane, and increased levels of KChol are associated with numerous pathologies. It is thought to induce apoptosis via inactivation of the phosphatidylinositol 3-kinase/Akt signaling pathway — a pathway that depends on lipid-rafts as signaling platforms. By means of coarse-grained molecular dynamics simulations, we demonstrate that KChol disrupts the liquid-liquid phase separation seen in an equimolar mixture of (dipalmitoylphosphatidylcholine) DPPC, (dioleoylphosphatidylcholine) DOPC, and Cholesterol (Chol). This disruption occurs via two mechanisms: i) KChol adopts a wider range of orientations with the membrane, which disrupts the packing of neighboring lipids and ii) KChol has no preference for DPPC over DOPC, which is the main driving force for lateral demixing in DPPC/DOPC/Chol membranes. This provides a molecular description of the means by which KChol induces apoptosis, and illustrates that a single chemical substitution to cholesterol can have a profound impact on the lateral organization of lipid membranes.

**Graphical TOC Entry:** 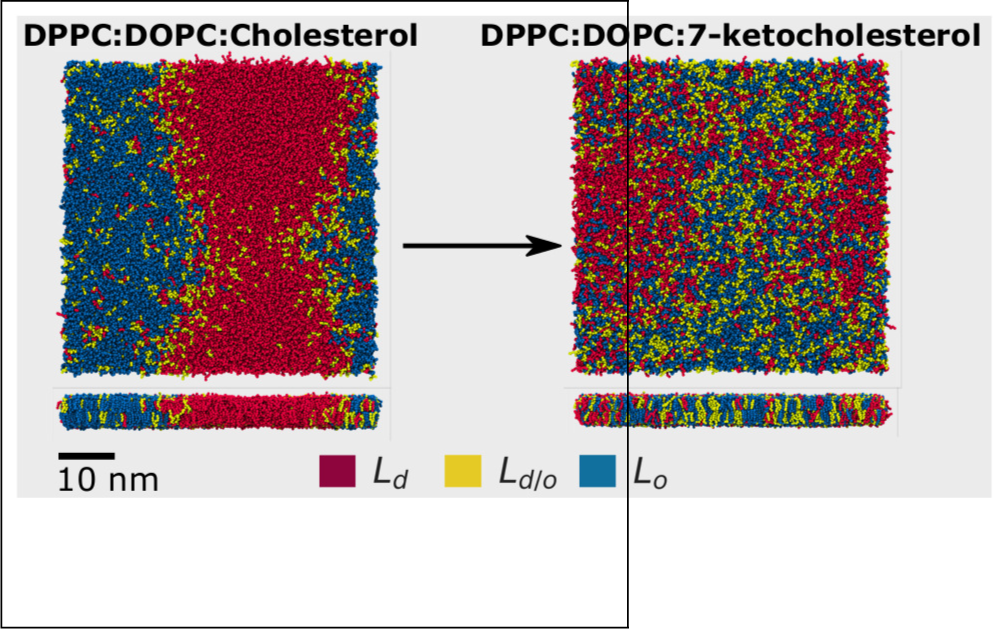

---

Since Simons and Ikonen first described lipid rafts,^1^ the existence, origin and nature of these structures in cellular membranes has been hotly debated.^2–17^ However, there is now direct evidence of microdomains in live yeast cell organelles;^18,19^ of nanodomains in live plant cell plasma membranes;^20^ of functional membrane microdomains in live bacteria;^21,22^ and of nanodomains in isolated mammalian cell plasma membranes.^23^ The ubiquitous presence of lipid-raft-like structures across the domains of life means their biological significance is no longer in doubt. This is further emphasized by their suspected roles in many membrane processes: from membrane signaling^24^ to membrane trafficking,^25^ from membrane deformation^26^ to membrane vesiculation,^27^ and from sites for oligomerization of peptides^28^ to sites for attachment of pathogens.^29^

Given the biological importance of lipid rafts, the disruption of liquid-ordered domains has the potential to impact myriad biological pathways and processes. Elevated levels of ring-oxidised sterols — produced by the autoxidation of cholesterol^30^ — are implicated in numerous pathologies,^31–43^ and have been speculated to prevent liquid-ordered domain formation.^44–46^ 7-ketocholesterol (KChol; Figure 1A) is one of the most abundant and cytotoxic oxysterols,^44^ and its presence in lipid rafts can induce cell death.^33^ KChol causes apoptosis via inactivation of the phosphatidylinositol 3-kinase/Akt signaling pathway^47^ — a pathway that depends on lipid rafts as signaling platforms.^24^ Further, by excluding KChol from lipid rafts, cell death is avoided.^48^ It is therefore possible that KChol induces apoptosis via disrupting the formation of liquid-ordered domains in the plasma membrane.

**Figure 1:**
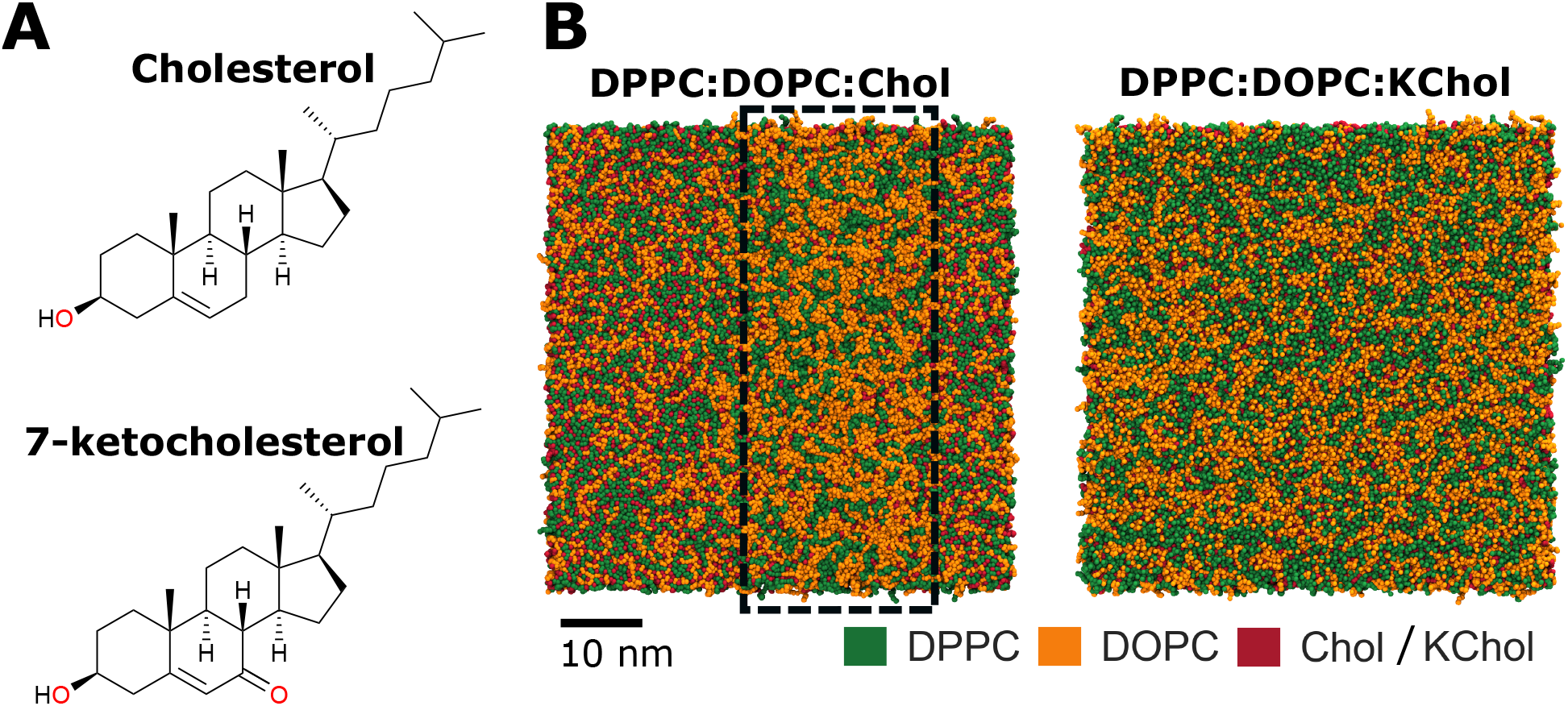
(A) Chemical structure of cholesterol and 7-ketocholesterol. (B) Lateral lipid distribution at 20 µs. The column highlighted in the DPPC:DOPC:Chol mixture is depleted of cholesterol.

The concept of lipid rafts originated as an explanation for the dynamic clustering of cholesterol (Chol; Figure 1A) and sphingolipids in the plasma membrane, and the preferential sorting of certain proteins into these domains.^1^ Since then, many different lipid mixtures have been found to be capable of nano- or micro-domain formation. ^17,49–56^ Indeed, the plasma membrane is thought to consist of many different raft-like and non-raft-like regions of varying lipid composition.^6,10,53^ These raft-like regions may arise through many different physical processes,^9,57,58^ with different physical mechanisms dominating at different stages of domain formation.^59^ Given the complexity of the plasma membrane, model membranes are typically employed for the study of domain formation. Membranes consisting of 1,2-dipalmitoyl-sn-3-phosphocholine (DPPC), 1,2-dioleoyl-sn-glycero-3-phosphocholine (DOPC) and Chol were the first phase-separating ternary mixture to have its phase boundaries fully mapped,^49^ and has since become the canonical mixture for studying phase separation in lipid membranes. While this mixture produces macroscopic phase separation, nanodomains behave surprisingly like genuine phases and so studying macroscopic phase separation may also inform us about nanodomains and lipid rafts.^60^

In this work, we report on the effect of cholesterol oxidation on domain formation studied by means of coarse-grained molecular dynamics simulations. We have studied equimolar mixtures of DPPC:DOPC:Chol and DPPC:DOPC:KChol using the MARTINI 2 force field with the Melo et al. cholesterol parameters.^61**?**^ We used the CHARMM-GUI MARTINI Maker^62,63^ to construct large bilayers (with 6,000 lipids per leaflet). At this size, the resultant phase-separated domains should be equivalent to those at the thermodynamic limit.^64^ We ran two replicas of each mixture for 20 µs at a temperature of 310 K.

We see a lateral demixing of lipids in the DPPC:DOPC:Chol membrane, with clearly defined Chol-poor and Chol-enriched regions (Figure 1B). On the other hand, lipids in the DPPC:DOPC:KChol membrane appear more uniformly distributed (Figure 1B). This immediately illustrates the profound impact that a single chemical substitution within one of the lipid constituents has on the lipid mixing within the membrane. To quantify the demixing of lipids in the membranes, we calculated the lipid enrichment/depletion index^65,66^ of each species over the final 4 µs of simulation time (Figure 2A). We see that Chol has a clear preference for DPPC over DOPC, and that DOPC tends to self-aggregate. This affinity between Chol and DPPC is what drives the macroscopic phase separation in the DPPC:DOPC:Chol membrane.^59,67^ Several recent studies have shown that small changes in a lipid’s chemistry can alter its affinity for Chol, and thus change the size and stability of lateral heterogeneities.^52,56,68–72^ Here we see that KChol, an oxidation product of Chol, has significantly less affinity for DPPC over DOPC than Chol — and the result is a disruption of the macroscopic phase separation.

**Figure 2:**
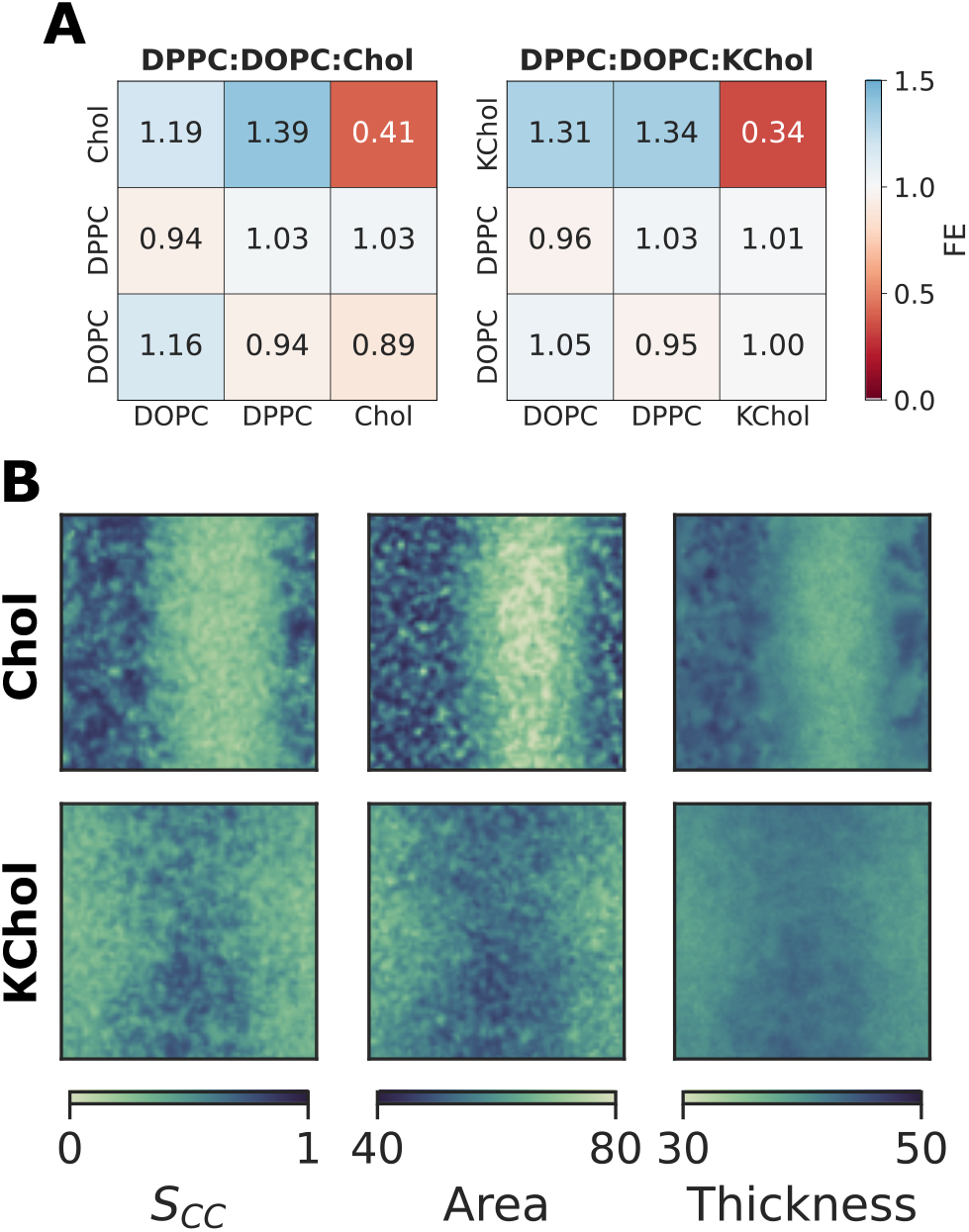
(A) Fractional enrichment of lipid species, calculated using the final 4 µs of each simulation. Values above and below 1 indicate enrichment and depletion, respectively. (B) Projection onto the membrane plane of the coarse-grained order parameter (*S*_*CC*_), area per lipid (Å^2^), and local membrane thickness (Å) of the phospholipids.

As a result of the phase separation, we see a large order gradient across the DPPC:DOPC:Chol membrane — the Chol-depleted region in Figure 1B is significantly more disordered than the Chol-enriched region. It has a larger area per lipid, smaller membrane thickness, and more disordered acyl tails than the Chol-enriched region (Figure 2B, upper panel). Whilst there is little lateral demixing of lipid species in the DPPC:DOPC:KChol membrane, we still observe an order gradient across the membrane (Figure 2B, lower panel). There is a large ordered region in the center of this membrane, with disordered regions either side. There is, however, a reduced gradient compared to the one in DPPC:DOPC:Chol membrane, and the boundary between the ordered and disordered domains is more diffuse. The lateral heterogenity in the DPPC:DOPC:KChol membrane is thus more akin to the nanodomains that form in DPPC:cholesterol binary mixtures^73^ than to the phase separated DPPC:DOPC:Chol membrane.

To better understand the affect of cholesterol oxidation on the domain-formation process, we constructed hidden Markov models (HMM) based on lipid thicknesses to assign each lipid molecule at each frame to one of three states: ordered (*L*_*o*_), disordered (*L*_*d*_), or intermediate (*L*_*d/o*_). We generated HMMs based on the methodology proposed by Park and Im.^74^ Briefly, we first calculated the thickness of each phospholipid molecule as the mean thickness in *z* of its two acyl tails, and the thickness of each sterol as the extent in *z* of the entire molecule. Then, for each lipid species, we binned these thicknesses into nine states, which served as the emission states of the model. We used a Gaussian mixture model to initialize the parameters (*µ, σ*) of the hidden Gaussian distributions, before using the Baum-Welch algorithm to fit the model parameters based on the emission states and initial parameters. Finally, we used the Viterbi algorithm to decode the most likely time series of hidden states (*L*_*o*_, *L*_*d*_, or *L*_*d/o*_) for each lipid.

The lateral distribution of ordered states can be seen in Figure 3. At 20 µs, the *L*_*o*_ and *L*_*d*_ regions of the DPPC:DOPC:Chol membrane clearly correspond to the ordered and disordered regions, respectively, seen in Figure 2B. Further, the *L*_*o/d*_ lipids are predominantly found at the *L*_*o*_-*L*_*d*_ interface, giving us confidence that the HMM has accurately assigned lipids to the correct ordered state.

**Figure 3:**
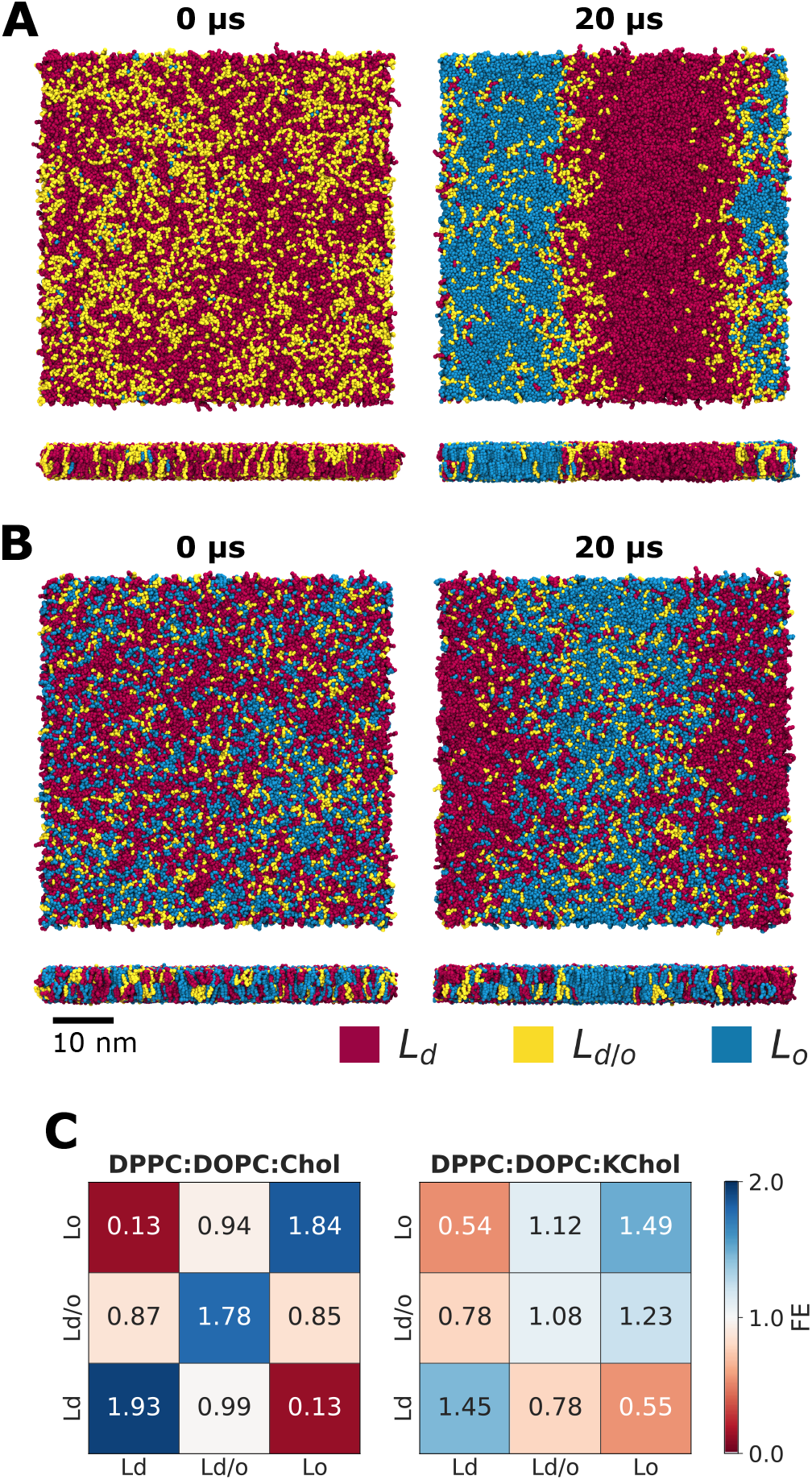
Lateral distribution of ordered (*L*_*o*_), disordered (*L*_*d*_), and intermediate (*L*_*d/o*_) lipids in the (A) DPPC:DOPC:Chol membrane and (B) DPPC:DOPC:KChol membrane. (C) Fractional enrichment of lipids by their phase (*L*_*d*_, *L*_*d/o*_, *L*_*o*_), calculated using the final 4 µs of each simulation. Values above and below 1 indicate enrichment and depletion, respectively.

At 0 µs, very few lipids in the DPPC:DOPC:Chol membrane are in the *L*_*o*_ state — almost all lipids are either *L*_*d*_ or *L*_*d/o*_ (Figure 3A). In particular, cholesterol is mostly *L*_*d*_ whereas the phospholipids are predominantly in the intermediate *L*_*d/o*_ state (Figure S2C). These *L*_*d*_ and *L*_*d/o*_ lipids are initially evenly distributed within the bilayer, with no sign of *L*_*o*_ domains. The domain formation process begins with a demixing of the *L*_*d*_ and *L*_*d/o*_ lipids (Figure S2A). *L*_*d*_ Chol then proceeds to become more ordered (Figure S2C). This in turn causes the DPPC and DOPC molecules in the intermediate state to also transition into the ordered *L*_*o*_ state. This transition from *L*_*d*_ and *L*_*d/o*_ to *L*_*o*_ is almost complete by 5 µs. Over time, the boundary between the *L*_*o*_ and *L*_*d*_ regions becomes more well-defined, and the phase separation is nearly complete by 10 µs (Figure S2A). By 20 µs, there are two clear phases present in the DPPC:DOPC:Chol membrane, and there is a significant enrichment of *L*_*o*_ lipids around other *L*_*o*_ lipids (Figure 3C).

Conversely, in the DPPC:DOPC:KChol membrane there is little change in the fraction of lipids in *L*_*o*_ state over time (Figure S2D). Instead, the *L*_*d*_ and *L*_*o*_ regions seen in Figure 3B form via a lateral demixing of the ordered and disordered lipids. Unlike in the DPPC:DOPC:Chol membrane, this demixing is not followed by an increased ordering of the *L*_*o*_ acyl tails (Figure S2B). In fact, there is very little change in the ordering of acyl tails in the DPPC:DOPC:KChol membrane throughout the simulation (Figure S3). This is in clear contrast to the acyl tails in the DPPC:DOPC:Chol membrane, which become significantly more ordered. The result is two co-existing macroscopic phases in the DPPC:DOPC:Chol membrane, but smaller, less stable nanodomains in the DPPC:DOPC:KChol membrane.

We see that the *L*_*o*_ and *L*_*d*_ regions in the DPPC:DOPC:KChol membrane are much more alike than those of the DPPC:DOPC:Chol mixture (Table 1). Generally, the *L*_*d*_ lipids of the KChol membrane are less disordered than those of the Chol membrane, while the *L*_*o*_ lipids of the KChol membrane are less ordered than those of the Chol membrane. We do, however, see that the *L*_*d*_ lipids in the KChol membrane have a larger area per lipid than the *L*_*d*_ lipid molecules in the Chol membrane, albeit only by 0.2 Å. This is likely because KChol adopts a wider range of orientations in the membrane (Figure S4). Chol has a strong tendency to be oriented at around 10° (and 170°), whereas KChol has a broader distribution of orientations with a peak at around 15° (and 165°). KChol adopts a wider range of orientations in the membrane so that its hydrophilic ketone group can be exposed to the solvent. This increased orientational freedom of KChol will likely lead to an increased area per lipid. An implication of this is that KChol will disrupt the local packing of lipids in the *L*_*o*_ phase.^44,46^ This therefore explains why the order gradient in the DPPC:DOPC:KChol membrane does not increase after the demixing of *L*_*o*_ and *L*_*d*_ lipids, unlike in the DPPC:DOPC:Chol membrane.

**Table 1:** Mean area per lipid, coarse-grained order parameter (*S*_*CC*_), and membrane thickness over the final 4 µs of simulation time. The standard error is 0.01 or less for all values.

|  | System | Lipid |  |  |  |  |  |
| --- | --- | --- | --- | --- | --- | --- | --- |
|  |  | DPPC |  | DOPC |  | Chol/KChol |  |
| | | $L_d$ | $L_o$ | $L_d$ | $L_o$ | $L_d$ | $L_o$ |
| Area ($\text{\AA}^2$) | DPPC:DOPC:Chol | 66.5 | 50.2 | 69.3 | 51.1 | 35.4 | 29.8 |
|  | DPPC:DOPC:KChol | 59.6 | 52.6 | 62.8 | 55.7 | 35.6 | 32.0 |
| $S_{CC}$ | DPPC:DOPC:Chol | 0.38 | 0.75 | 0.28 | 0.65 | - | - |
|  | DPPC:DOPC:KChol | 0.48 | 0.68 | 0.31 | 0.55 | - | - |
| Thickness ($\text{\AA}$) | DPPC:DOPC:Chol | 39.4 | 43.9 | 39.1 | 43.7 | - | - |
|  | DPPC:DOPC:KChol | 40.7 | 42.6 | 40.3 | 41.8 | - | - |

Within the two mixtures, we see a difference in the lipid composition of their respective *L*_*o*_ domains (Figure S5). The *L*_*o*_ domain of the DPPC:DOPC:KChol membrane is enriched in DPPC (with a DPPC:DOPC:KChol ratio of 0.38 : 0.29 : 0.33), and we observe no significant change in its composition over the course of 20 µs. On the other hand, in the DPPC:DOPC:Chol membrane, we observe small ordered clusters enriched in Chol forming at the beginning of the simulation. Then, we find that the onset of nanodomain formation is associated with an increase in other lipid species, especially DPPC, in the Chol-enriched ordered clusters. Despite the other species joining the *L*_*o*_ domain, it remains enriched in Chol even at 20 µs (with a DPPC:DOPC:Chol ratio of 0.34 : 0.22 : 0.44).

The resulting *L*_*o*_ domain of DPPC:DOPC:Chol is not only more ordered than that of DPPC:DOPC:KChol, it is also larger and more stable. Figure 4A shows the largest cluster of *L*_*o*_ lipids in the upper leaflet of each mixture over time. From around 2 µs onward, almost all *L*_*o*_ lipids in the DPPC:DOPC:Chol mixture are part of the *L*_*o*_ domain. Conversely, the largest *L*_*o*_ cluster in the DPPC:DOPC:KChol membrane dissociates at around 5 µs before reforming, and even at 20 µs no more than 80% of *L*_*o*_ lipids are in the *L*_*o*_ domain. This dissociation of the *L*_*o*_ domain coincides with a decrease in the interleaflet registration of the *L*_*o*_ domains (Figure 4B). Domain registration in the Chol membrane, however, is also faster and more stable.

**Figure 4:**
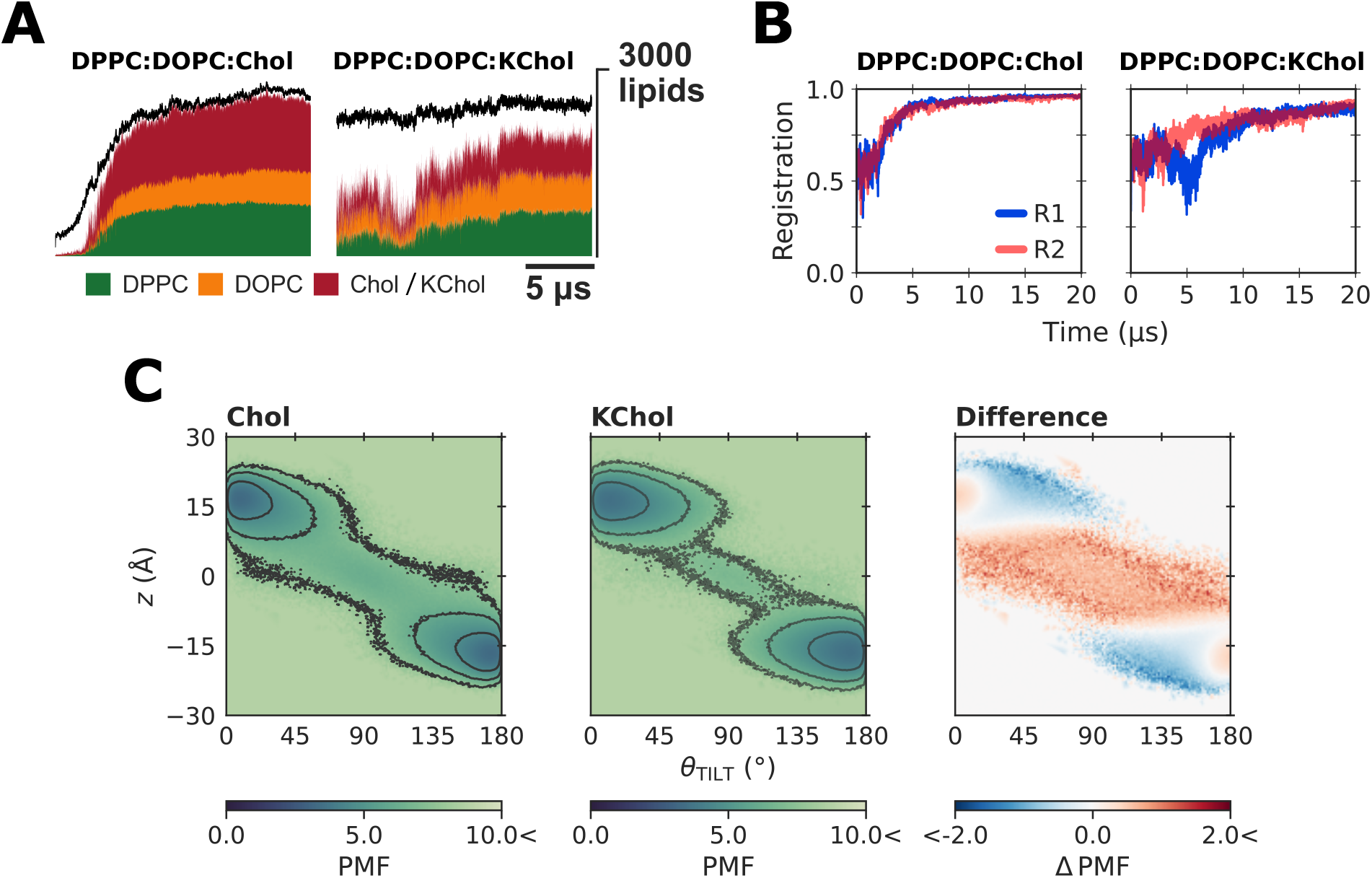
(A) Number of each lipid species in the largest cluster of *L*_*o*_ lipids. The black curve shows the total number of *L*_*o*_ lipids present. (B) Domain registration. (C) Potential of Mean Force (PMF) of sterol orientation (*θ*_TILT_) and height (*z*). For Chol, there is a free energy barrier of around 5 kcal mol^−1^ in the region −12 Å *< z <* 12 Å, 65° *< θ*_TILT_ *<* 115°. The difference plot shows PMF_KChol_ − PMF_Chol_; red regions are less favorable for Kchol and blue regions more favorable.

It is not only the structure of the *L*_*o*_ domains that changes upon oxidation, but also the dynamics of the molecules within the domains. In the DPPC:DOPC:Chol mixture, the lateral diffusion of *L*_*o*_-domain lipids is 1.5 times faster than those in the largest cluster of *L*_*d*_ lipids (Table 2), which is in line with atomistic simulations and experimental measurements.^66^ This difference is significantly reduced in the DPPC:DOPC:KChol membrane (Table 2) — a result of the fact in this membrane we see nanodomain formation, with a smaller order gradient, rather than microdomain formation, with a larger order gradient.^75^

**Table 2:** Flip-flop rate, *k* (×10^6^ s^−1^), of cholesterol and 7-ketocholesterol, and lateral diffusion coefficient, *D*_*xy*_ (×10^−7^ cm^2^ s^−1^), of lipids in the *L*_*o*_ and *L*_*d*_ domains. Values calculated using the final 4 µs of each trajectory.

| | $k$ | $D_{xy}$ | |
| --- | --- | --- | --- |
| | | $L_o$ | $L_d$ |
| Chol | 1.80 | 3.4 | 5.1 |
| KChol | 0.64 | 3.2 | 3.7 |

We also find a substantial affect on interleaflet dynamics upon cholesterol oxidation. The rate of cholesterol flip-flop in the DPPC:DOPC:Chol membrane is around 3 times faster than in the DPPC:DOPC:KChol membrane (Table 2). The reduced flip-flop rate for KChol is the result of an increased free energy barrier to translocation compared to Chol. The potentials of mean force (PMF) for the height and orientation of Chol and KChol within the membranes are shown in Figure 4C. For Chol, there is a barrier to flip-flop of around 5 kcal mol^−1^. This is due to the unfavorable desolvation of the hydroxyl group during the flipflop process, which occurs as the sterol crosses through the hydrophobic core of the bilayer (−12 Å*< z <* 12 Å) and rotates to align with lipids in the apposing leaflet (65° *< θ*_TILT_ *<* 115°). The ring-oxidation of cholesterol into 7-ketocholesterol further increases this barrier by another 2 kcal mol^−1^ (Figure 4C). This is because both the ketone and hydroxyl groups must be desolvated for flip-flop to occur. The result is that KChol is less likely move to the midplane and we thus see a reduced rate of flip-flop in the DPPC:DOPC:KChol mixture.

We have shown that the macroscopic phase separation seen in a DPPC:DOPC:Chol membrane is disrupted by the autoxidation of cholesterol into 7-ketocholesterol. In a DPPC:DOPC:Kchol membrane, we instead see nanodomain formation that is more akin to that expected in the plasma membrane.^6,10,15,53^ This disruption arises from the hydrophilicity of the ketone group of KChol, which has two effects on the domain formation. First, to allow for the hydration of the ketone group, KChol adopts a broader distribution of orientations in the membrane. This disrupts the local packing of lipids, inducing disorder in *L*_*o*_ regions.^44–46^ Second, the reason Chol prefers to interact with DPPC over DOPC is because DPPC is better at shielding the hydrophobic rings of cholesterol from the surrounding solvent. The tetracyclic rings of KChol, however, are less hydrophobic due to the presence of the ketone group; KChol tends to expose this moiety to the solvent rather than seeking refuge in the hydrophobic core of the bilayer, and we thus find that KChol has less of a preference for DPPC over DOPC. This reduced preference for DPPC suppresses the lateral demixing of lipids, which in turn disrupts the liquid-liquid phase separation seen in the DPPC:DOPC:Chol mixture. We also see that the hydrophilicity of the KChol ketone group suppresses translocation. The reduced rate of translocation has little effect on domain registration at in the equilibrated mixture studied here, but in more physiologically-relevant mixtures sterol flip-flop is required for interleaflet domain registration.^54^

Chol preferentially mixes with sphingolipids over glycerophospholipids for the same reason it prefers DPPC over DOPC - sphingolipids are better at shielding cholesterol from the surrounding solvent.^76^ We can therefore expect the increased hydrophilicity of KChol to diminish its affinity for sphingolipids compared to cholesterol. This would either disrupt the formation of Chol-sphingolipid nanodomains in biological membranes, or at least reduce the lateral order gradient as seen here. The reduced order gradient would have implications lipid-raft protein-sorting due to the hydrophobic mismatch between raft regions and their embedded proteins. Such implications include the disruption of cell-signaling pathways, via which KChol is known to induce apoptosis.

## Methods

We used the CHARMM-GUI MARTINI Maker^62,63^ to construct an equimolar mixture of DPPC:DOPC:Chol with 6,000 lipids per leaflet. The system had 89,995 non-polarizable water beads (10% of which were anti-freeze beads), 1,154 Na beads, and 1,154 Cl beads. We used the MARTINI 2 force field along with the Melo et al. parameters for cholesterol.^61,77^ To construct the DPPC:DOPC:KChol membrane, we used the DPPC:DOPC:Chol bilayer and replaced all Chol molecules with KChol molecules. We first performed a steepest descent minimization for 5,000 steps in which the sterol constraints were replaced by harmonic bonds. We then performed a series of short (*∼*1 ns) equilibrations, with increasing timesteps (2, 5, 10, 15, 20 fs), to relax the systems. In these equlibrations, we applied position restraints with decreasing coefficients (200, 100, 50, 20, 10 kcal mol^−1^) in the *z*-dimension to the PO4 and ROH beads. For production simulations, we used a timestep of 25 fs and to suppress large-scale undulations^78^ we applied a 2 kcal mol^−1^ restraint in the *z*-dimension to the PO4 beads of phospholipids in the upper leaflet. All simulations were performed using the semi-isotropic NPT ensemble at 310K and 1 bar, and using the new-RF parameter set^79^ for performing MARTINI simulations with GROMACS. To perform the second replicas of each mixture we used a different random seed when generating initial velocities. Coordinates were stored every 0.25 ns. All simulations were performed using GROMACS 2018.2. ^80^

## Acknowledgement

Via our membership of the UK’s HEC Materials Chemistry Consortium, which is funded by EPSRC (EP/L000202/1, EP/R029431/1), this work used the ARCHER UK National Supercomputing Service (http://www.archer.ac.uk) and the UK Materials and Molecular Modelling Hub (MMM Hub) for computational resources, which is partially funded by EPSRC (EP/P020194/1) to carry out the MD simulations reported in this manuscript. P.S. acknowledges the funding provided by the EPSRC DTP Studentship Block Grant (EP/N509498/1). The idea for this work was generated by discussions which were had during the EPSRC-funded project “Red Cell Physical Properties in Health and Disease” (EP/N007700/1). C.D.L. acknowledges the supportive research environment of the EPSRC Center for Doctoral Training in Cross-Disciplinary Approaches to Non-Equilibrium Systems (CANES, No. EP/L015854/1).

## Supporting Information Available

Parameters for a MARTINI model of 7-ketocholesterol. Detailed analysis methods. Snapshots illustrating the dynamics of domain formation. Distribution of the coarse-grained order parameter. Fractional composition of the largest *L*_*o*_ domain. Distribution of sterol orientation.

## Notes

### Competing Interest Statement

The authors have declared no competing interest.

### Summary of Updates

Clarification of the force field used for the simulations.

