## Supporting Information for "Cholesterol Oxidation Modulates the Formation of Liquid-Ordered Domains in Model Membranes"


Paul Smith,[*,†] Peter G. Petrov,[‡] and Christian D. Lorenz[*,†]

†*Department of Physics, King's College London, London, WC2R 2LS, UK*

‡*Department of Physics and Astronomy, University of Exeter, Stocker Road, Exeter EX4 4QL, UK*

**Listing S1**: Parameters for a MARTINI model of 7-ketocholesterol

**Figure S1**: Structure of the MARTINI model of Chol/KChol

**Figure S2**: Lateral distribution of ordered states

**Figure S3**: Distribution of the coarse-grained order parameter

**Figure S4**: Distribution of sterol orientations

**Figure S5**: Fractional composition of the $L_o$ domain

### MARTINI parameters for 7-ketocholesterol

In keeping with the modular philosophy of MARTINI,[1] we modeled 7-ketocholesterol by changing the R2 bead type from SC3 (semi-repulsive to water) to SN0 (intermediate with water), which has the same mass as SC3 but is more polar. We experimented with using N0 instead of SN0, but the non-bonded interactions with the solvent were too strong and led to freezing of water at the bilayer interface, even with 10% anti-freeze water beads.

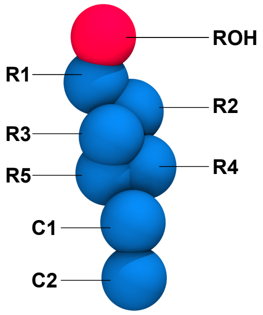


Figure S1: Structure of the MARTINI model of Chol/KChol. The R2 bead is of type SC3 and SN0 for Chol and KChol respectively. The red ROH bead is polar, representing the hydroxyl group of the sterols, and the blue beads are apolar.

SN0 is a conservative choice of bead for the ketone group. The SP1 bead (polar; almost attractive with water) is used for the ROH group in the MARTINI model of cholesterol, as well as in all three ROH groups of the cholate MARTINI model.[2] These hydroxyl groups, however, are more polar than the ketone group of KChol — hydroxyl moeities can both donate and accept hydrogen bonds. We therefore used the SN0 bead (intermediate polar; intermediate with water). The mass of the beads and all bond lengths are unchanged, which is in keeping with the cholate MARTINI model (the ROH beads in cholate keep the same mass as the corresponding beads in the cholesterol model).

The angular orientation of our KChol model is in line with previous atomistic simulations of this ring-oxidized sterol.[3–5] This is an important differentiator between Chol and KChol, thus giving confidence in the KChol model.

```
[ moleculetype ]
; molname  nrexcl
  KCHL     1

[ atoms ]
; i type resnr residue atom cgnr charge mass
  1 SP1  1     KCHL    ROH  1    0.0    77.22
  2 SC1  1     KCHL    R1   2    0.0      0.0
  3 SN0  1     KCHL    R2   3    0.0    38.69
  4 SC1  1     KCHL    R3   4    0.0   159.65
  5 SC1  1     KCHL    R4   5    0.0      0.0
  6 SC1  1     KCHL    R5   6    0.0      0.0
  7 SC1  1     KCHL    C1   7    0.0    39.44
  8  C1  1     KCHL    C2   8    0.0     72.0

[ bonds ]
; i  j  funct  length  force
  7  8  1      0.425   1250.0
#ifndef FLEXIBLE
[ constraints ]
#endif
  1  3  1    0.4904  1000000
  1  4  1    0.6019  1000000
  3  4  1    0.2719  1000000
  7  3  1    0.7237  1000000
  7  4  1    0.5376  1000000

[ dihedrals ]
  1  3  4  7   2  -179.7  50

[ virtual_sites3 ]
; In-plane bead from frame 4-3-7 (bead 5)
  5  4  3  7   1  0.9613  0.6320
```

```
; Out-of-plane bead from frame 3-1-4 (bead 2)
  2  3  1  4   4  0.5207  0.2882  -0.83824
; Out-of-plane bead from frame 4-3-7 (bead 6)
  6  4  3  7   4  0.2287  0.4111   1.1531

[ angles ]
; i  j  k  funct  angle  force
  4  7  8  2      180.0  25.0

[ exclusions ]
; i  j  k  ...
  1  2  3  4  5  6  7
  2  3  4  5  6  7
  3  4  5  6  7
  4  5  6  7
  5  6  7
  6  7

#ifdef BILAYER_LIPIDHEAD_FC
    [ position_restraints ]
    ;to fix Z postion of head grop in bilayer simulation
      1     1.0      0.0      0.0     BILAYER_LIPIDHEAD_FC
#endif
```

Listing 1: Parameters for a MARTINI model of 7-ketocholesterol. Adapted from the equilibrium structure of the Melo et al.[2] cholesterol model for MARTINI. Replaces the SC1 bead at R2 with an SN0 bead to reflect the increased polarity after oxidation.

### Analysis methods

Analysis was performed using MDAnalysis,[6,7] LiPyphilic,[8] FATSLiM,[9] SciPy,[10] and HMMLearn.[11] Unless stated otherwise, every tenth frame (5 ns) was used in the analysis. The standard errors reported in Table 1 were calculated using 50 ns block averages.

***Hidden Markov Model*** Lipids were assigned to be either ordered ($L_o$), disordered ($L_d$), or intermediate ($L_{d/o}$) by constructing Hidden Markov Models, as described in the main text. SciPy[10] was used to generate the Gaussian Mixture Model, from which the initial HMM parameters were derived. HMMLearn was then used to refine the model parameters and subsequently decode the most likely sequence of ordered states. Smith et al.[12] describes this procedure in more detail.

***Area per lipid*** The area per lipid was calculated via a Voronoi tessellation of the $x$ and $y$ coordinates of GL1, GL2, and ROH beads within each leaflet. The analysis was performed using LiPyphlic,[8] which uses Freud[13] to perform the tessellation of atomic coordinates.

***Coarse-grained order parameter*** The coarse-grained order parameter, $S_{CC}$, is given by:

$$S_{CC} = \frac{\langle 3\cos^2\theta \rangle}{2}$$

where $\theta$ is the angle between the membrane normal (approximated as the $z$-axis) and the vector connecting two consecutive tail beads. The average is taken over all beads in a molecule. LiPyphilic[8] was used to perform the calculation.

***Membrane thickness*** For each phospholipid, a local leaflet patch was defined by all PO4 beads within 60 Å of the reference lipid's PO4 bead. The normal to this patch was used to identify a reference lipid for the apposing leaflet, and a local patch defined for this second lipid in a similar manner. The membrane thickness for the original lipid was taken to be the distance between the center of mass of the two leaflet patches. FATSLiM[9] was used to perform the calculation.

***Fractional enrichment*** To calculate the fractional enrichment of lipid species, a neighbor matrix, $A$, was first constructed. The matrix is 12,000 by 12,000, where each row or column represents a distinct lipid molecule. $A_{ij} = 1$ if two lipids are neighbors and $A_{ij} = 0$ otherwise. Two lipids were considered neighbors is they have any of the GL1, GL2, or ROH beads within 15 Å of one another. The neighbor matrix was then used to determine the fractional enrichment of each species over the final 4 µs of simulation time. The fractional enrichment of species $B$ around species $A$, $E_{AB}$, is given by:

$$E_{AB} = \frac{[B]_{\text{Local}}}{[B]_{\text{Bulk}}}$$

where $[B]_{\text{Bulk}}$ and $[B]_{\text{Local}}$ are the bulk concentrations and local concentration around species A, respectively, of species B.[14]

The same neighbor matrix was used to calculate the fractional enrichment based on lipid order ($L_d$, $L_{d/o}$, or $L_o$) of lipids. LiPyphilic[8] was used to construct the neighbor matrix.

***Largest domain*** To calculate the largest cluster of $L_o$ lipids at a given frame, the neighbor matrix described above was used. First, the rows and columns of non-$L_o$ lipids were removed. Then the largest connected component of this new matrix was found, which corresponds to the largest cluster of $L_o$ lipids. The same approach was used to identify lipids in the largest $L_d$ domain at each frame. LiPyphilic[8] was used to find the largest clusters.

***Registration*** The interleaflet registration, $r_{u/l}$, can be defined as the Pearson correlation coefficient between lateral densities of $L_o$ lipids in the upper and lower leaflets.[15] Values of $r_{u/l} = 1$ correspond to perfectly registered domains and values of $r_{u/l} = -1$ correspond to perfectly anti-registered domains. LiPyphilic[8] was used to perform the calculation.

***Flip-flop*** To calculate the flip-flop rate of cholesterol, lipids were first assigned to leaflets based on the $z$ coordinate of their GL1, GL2, and ROH beads using LiPyphilic.[8] A cholesterol molecule with its ROH bead within 10 Å of its local membrane midpoint was classified as being in the midplane. A cholesterol molecule was taken to have flip-flopped if it left one leaflet, passed through the midplane, and the resided in the apposing leaflet for at least 10 ns.

Every tenth frame (0.5 ns) from the final 4 µs of each replica was used in the analysis. The rate was calculated by dividing the total number of observed flip-flops by the product of the number of sterol molecules and the total simulation time used for the analysis. LiPyphilic[8] was used to perform the calculation.

***PMF*** Sterol height was calculated as the signed distance in $z$ from the ROH bead to the membrane midpoint. Sterol orientation was defined as the angle between the $z$-axis and the vector from bead R5 to R1. The PMF of sterol orientation and height, $F(z, \theta_z)$, was then calculated directly from the joint probability distribution, $P(z, \theta_z)$. The PMF is given by:

$$F(z, \theta_z) = -k_B T \ln P(z, \theta_z)$$

where $k_B$ is the Boltzmann constant and $T$ is the temperature in Kelvin. LiPyphilic[8] was used to plot the PMFs.

***Lateral diffusion*** The lateral diffusion coefficient was calculated from the mean-square displacement (MSD) of PO4 and ROH beads via the Einstein relation. The MSD was calculated for lipids in the largest $L_o$ or $L_d$ clusters separately. The center of mass motion of the $L_o$ or $L_d$ cluster was removed from the MSD of the respective lipids. The MSD and diffusion coefficients were calculated using LiPyphilic,[8] which uses tidynamics[16] to calculate the MSD via the Fast Correlation Algorithm.

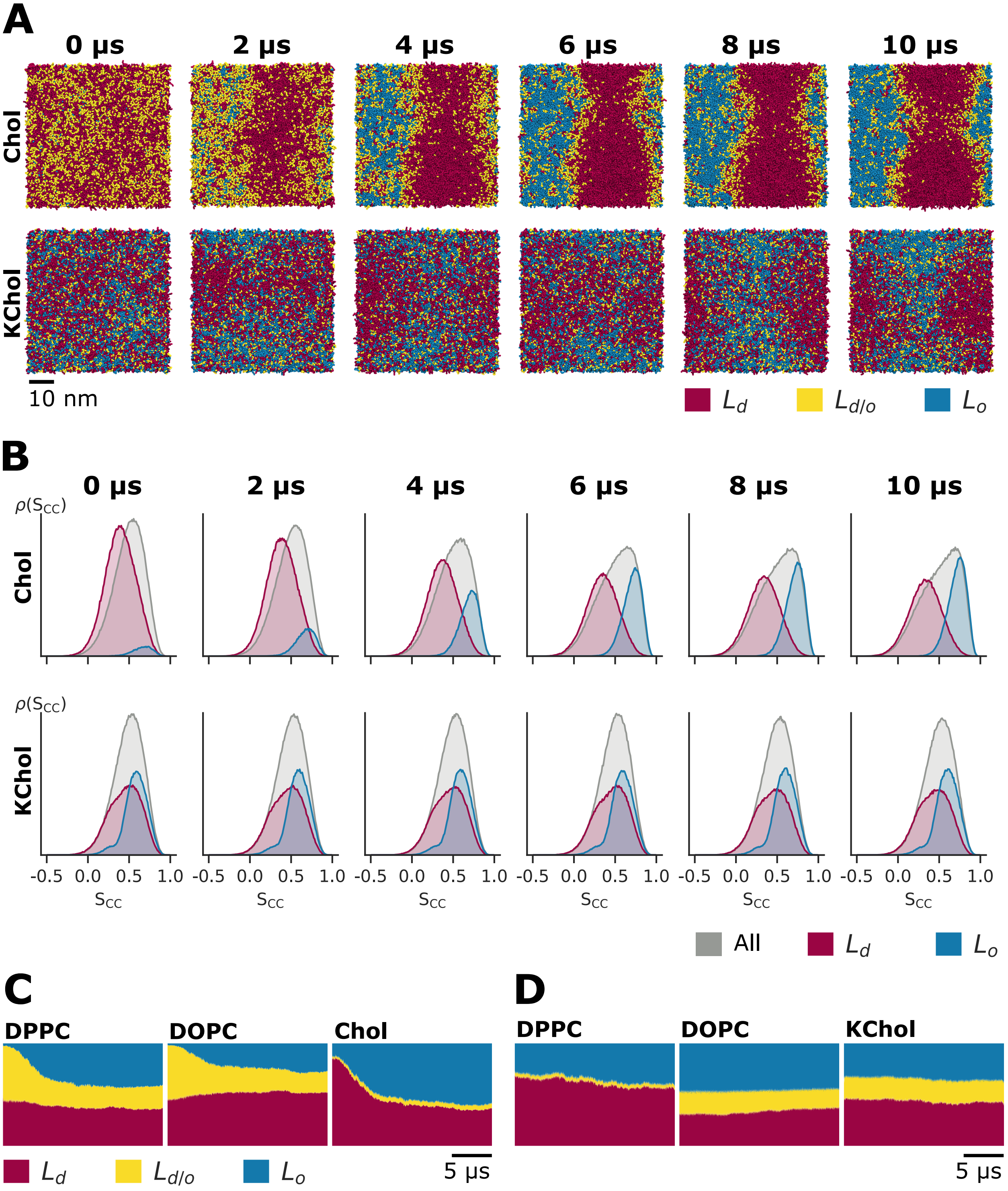


Figure S2: (A) Lateral distribution of $L_d$, $L_o$ and intermediate ($L_{d/o}$) lipids throughout the first 10 $\mu$s of simulation time. (B) Coarse-grained order parameter, $S_{CC}$, for phospholipids throughout the first 10 $\mu$s of simulation time. (C, D) Fraction of each lipid species in $L_d$, $L_o$ or intermediate states over time.

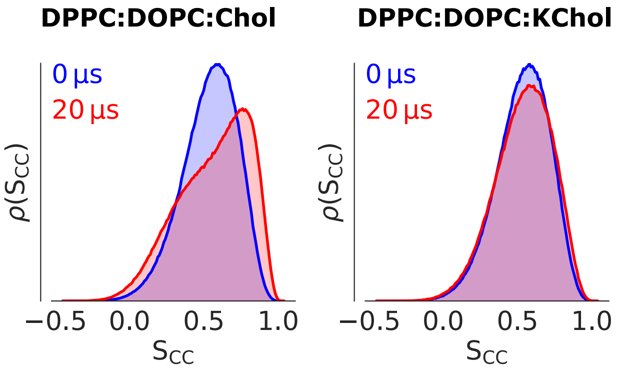


Figure S3: Coarse-grained order parameter, $S_{CC}$.

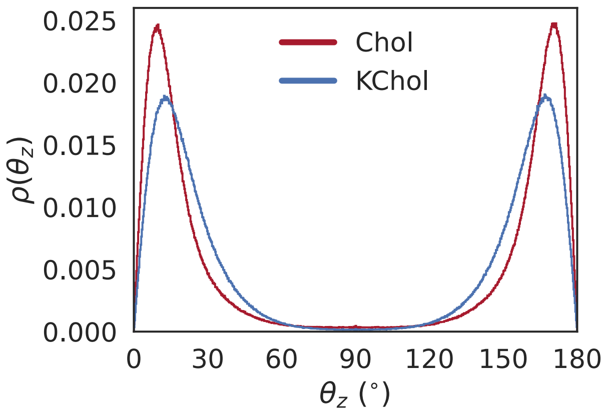


Figure S4: Sterol orientation, defined as the angle between the positive $z$ axis and the vector from bead R5 to bead R1.

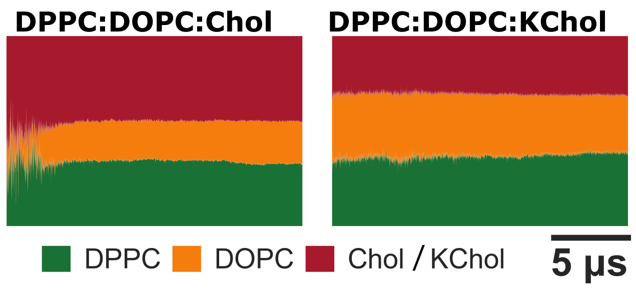


Figure S5: Fractional composition of the largest $L_o$ domain over time.
